# Reduction of LOXL2-mediated H3K4 oxidation increases chromatin accessibility and promotes chemosensitivity of triple-negative breast cancer cells

**DOI:** 10.1101/416495

**Authors:** J.P. Cebrià-Costa, L. Pascual-Reguant, A. Gonzalez-Perez, G. Serra-Bardenys, J. Querol, M. Cosín, G. Verde, R.A. Cigliano, W. Sanseverino, S. Segura-Bayona, A. Iturbide, D. Andreu, P. Nuciforo, C. Bernado-Morales, V. Rodilla, J. Arribas, J. Yelamos, A. Garcia de Herreros, T.H. Stracker, S. Peiró

## Abstract

Oxidation of H3 at lysine 4 (H3K4ox) by lysyl oxidase–like 2 (LOXL2) generates an H3 modification with an unknown physiological function. We find that LOXL2 and H3K4ox are higher in triple-negative breast cancer (TNBC) cell lines and patient–derived xenographs (PDXs) than those from other breast cancer subtypes. ChIP-seq revealed that H3K4ox is located primarily in heterochromatin, where it is involved in chromatin compaction. Knocking down LOXL2 reduces H3K4ox levels and causes chromatin decompaction, resulting in a sustained activation of the DNA damage response (DDR) and increased susceptibility to anticancer agents. This critical role that LOXL2 and oxidized H3 play in chromatin compaction and DDR suggests that functionally targeting LOXL2 could be a way to sensitize TNBC cells to conventional therapy.

## Introduction

Histone modifications contribute to gene regulation both by directly affecting chromatin structure and by recruiting effector proteins (1). Deregulation of this enzymatic system can contribute to disease, including cancer. The lysyl oxidase family of proteins are copper- and quinone-dependent amine oxidases that oxidize the amino group located in the epsilon-position in lysines, thereby generating an aldehyde group (2). One of the members of the LOX family, lysyl oxidase–like 2 (LOXL2), deaminates unmethylated and trimethylated lysine 4 in histone H3 (H3K4me3) through an amino-oxidase reaction that uses the Cu(II) ion and the internal co-factor lysine-tyrosylquinone (LTQ), releasing the amino group and converting K4 into an allysine (H3K4ox) (3,4). Generation of this peptidyl aldehyde likely alters the local macromolecular structure of chromatin and the nature of any protein-protein or protein–nucleic acid interactions. This is particularly relevant for gene regulation, as changes in the macromolecular status of histones can affect chromatin conformation (4-6).

LOXL2 is overexpressed in many tumors, and especially in breast cancers (7-9). In this light, it is intriguing that some breast cancers are intrinsically resistant to chemotherapy; for these subtypes, chemotherapy induces a mesenchymal phenotype through the epithelial-to-mesenchymal transition (EMT) (10). EMT is likely to be a critical switch for tumor cell invasiveness and cell death resistance (11-13) and to involve chromatin reorganization, as it requires dramatic changes in cellular characteristics and gene expression (6,14). Notably, the key transcription factor Snail1 interacts with LOXL2 (15), and LOXL2 H3K4 oxidase activity generates an H3K4ox that regulates the repression of the *E-cadherin* gene and heterochromatin transcription, which play roles in two essential steps of EMT (6,16).

Double-strand breaks (DSBs) are a major form of DNA damage and cause a signaling response that can activate cell cycle checkpoint arrest and cell fate decisions, such as apoptosis or senescence. Increasing evidence suggests that higher-order chromatin structure affects DSB repair and signaling (17). For example, the DNA damage response (DDR) actively regulates decondensation of chromatin after DSBs (18), and it is amplified when chromatin is in an “open” state (17). Similarly, DDR signaling is affected by chromatin compaction in a DNA damage–independent manner (19-22). Finally, cancer genome sequencing studies have shown substantial variation in somatic mutation rates, with increased rates in heterochromatin (closed chromatin) as compared in euchromatin (open chromatin) (23,24). Furthermore, DNA mismatch repair (MMR) is more efficient in euchromatic genomic regions than in heterochromatin, such that fewer mutations accumulate in these open genomic regions (25).

We addressed the physiological functions of H3K4ox using an in-house generated antibody specific for this modification to analyze the H3K4ox levels in distinct breast cancer subtypes. Intriguingly, both mesenchymal triple-negative breast cancer (TNBC) cell lines and breast cancer patient-derived xenographs (PDXs) had high H3K4ox levels that correlated with high LOXL2 expression, as compared to other subtypes. Using ChIP-seq to map its genome-wide localization, we found that H3K4ox was enriched in heterochromatin in TNBC cells, which are highly metastatic and chemotherapy resistant. Decreasing LOXL2 levels reduced the amount of H3K4ox in chromatin that results in a sustained activation of DDR and chromatin decondensation, suggesting that H3K4ox levels are crucial for maintaining chromatin structure. Further, both LOXL2 depletion and treatment of TNBC with chromatin-modifying drugs sensitized cancer cells to conventional treatments. Thus, targeting H3K4ox levels may open a new therapeutic window for this subtype of breast cancer.

## Results

### Generating an H3K4ox-specific antibody

We initially generated a specific antibody for the recently-discovered histone modification of H3K4ox, as a prerequisite for studying its physiological function. Since the aldehyde group generated by LOXL2 reaction on trimethylated lysine 4 is highly reactive, hence unfit for immunochemical studies, we hypothesized that a primary alcohol might provide a similarly oxygen-bearing yet less reactive functionality for generating a modification-specific antibody that provides a readout of H3K4ox (Fig. 1A). Hence a H3 peptide containing a 6-hydroxynorleucine residue as allysine isostere at position 4 was synthesized and used for rabbit immunization (Fig. 1B). The resulting H3K4ox antibody was specific for the H3K4ox peptide, and did not recognize peptides with unmodified H3, H3K9me3, or H3K4me3, in a wide range of experimental conditions (dot blots, Western blot, immunofluorescence, and ChIP experiments) (Fig. 1, C, D, and F). Immunostaining showed that the H3K4ox modification is located in the nuclei (Fig. 1D). Analysis of purified nucleosomes from 293T cells showed that H3K4ox levels increased, and H3K4me3 levels decreased, when nucleosomes were incubated with wild-type recombinant LOXL2 but not with a catalytically inactive LOXL2 (LOXL2mut) (3) (Fig. 1E, upper panel). Moreover, the levels of H3K4ox also increased in MCF7 cells transfected with LOXL2 as compared to cells transfected with the empty vector (Fig. 1E, lower panel). Finally, MDA-MB-231 cells infected with an shRNA targeting the human LOXL2 (LOXL2 KD) showed a specific reduction in H3K4ox levels as compared to cells infected with an irrelevant shRNA (Control), in both Western blots and in ChIP-PCR experiments using the *E-cadherin* gene promoter (CDH1) (Fig. 1F). Kinetics of the reaction using recombinant LOXL2 and nucleosomes revealed that levels of intermediate alcohol were maintained for 2 hours, after which they lowered (Fig. 1G). Thus, as the intermediate alcohol is relatively stable, the antibody we generated can be used as a readout of the oxidized histone H3K4.

**Fig. 1.**
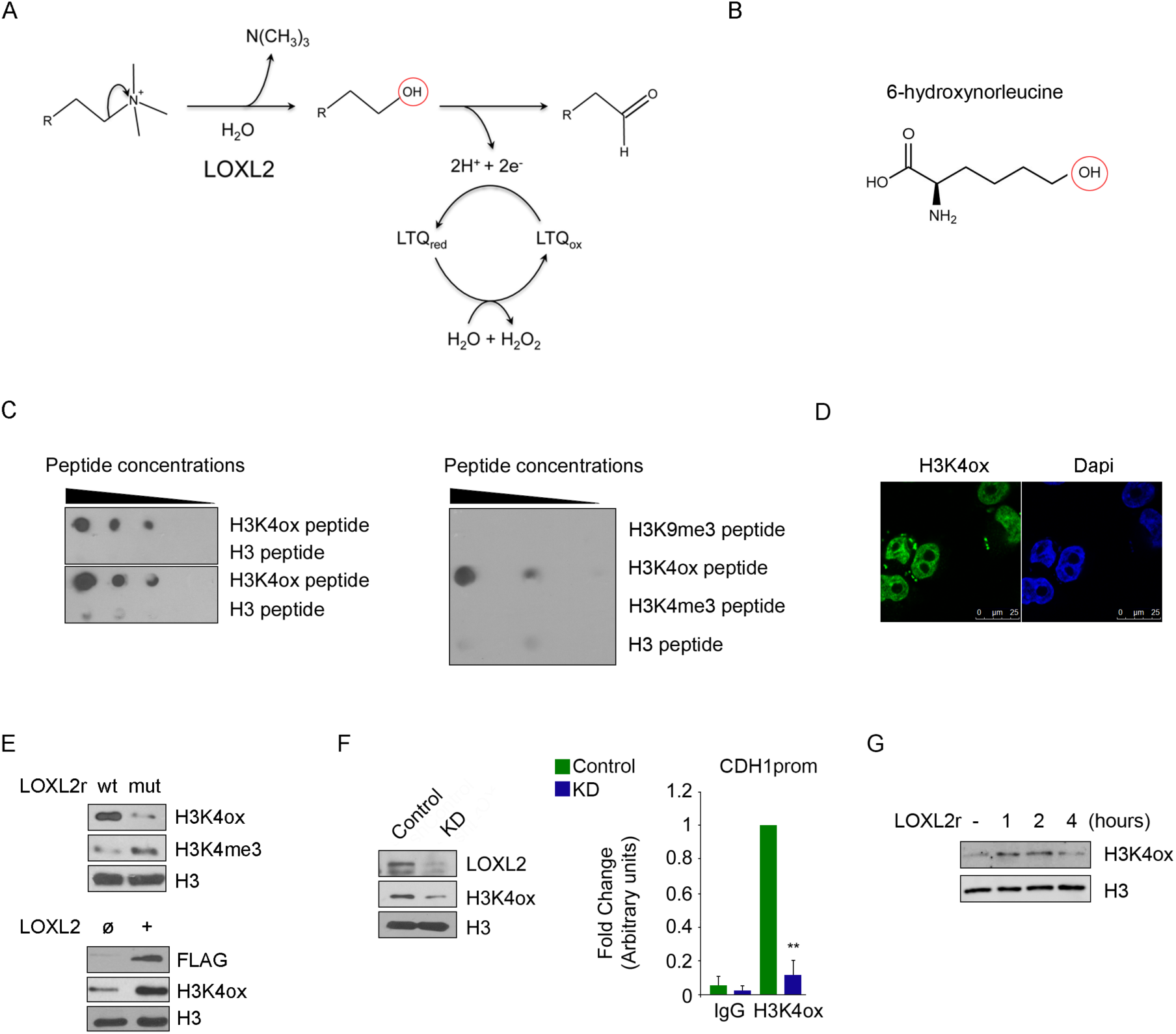
Quality control of the anti-H3K4ox antibody. (**A**) Schematic representation of the LOXL2 reaction. The red circle indicates the intermediate residue that is targeted by the anti-H3K4ox antibody. (**B**) The artificial amino acid 6-hydroxy-norleucine was used in the peptide to generate the anti-H3K4ox antibody. (**C**) The anti-H3K4ox antibody was specific in Western blot using in two replicates of dot blots of dilution series of oxidized histone H3 peptide (H3K4ox) or unmodified H3 peptide (left panel), as well as in a representative dot blot of a dilution series of H3K9me3, H3K4ox, H3K4me3, and H3 peptides (right panel). (**D**) The anti-H3K4ox antibody (green) was functional in confocal microscopy, as shown with MDA-MB-231 cells (green). Nuclei were staining with DAPI. (**E**) Nucleosomes were incubated with recombinant wild-type (wt) LOXL2 or a catalytically inactive LOXL2 (mut) (generated in baculovirus) to detect H3K4ox/H3K4me3 levels (upper panel). Lysates of MCF7 cells transfected with an empty vector (ø) or with LOXL2 were analyzed by Western blots, using the antibodies indicated (lower panel). (**F**) Western blot for LOXL2, H3K4ox, and total H3 in MDA-MB-231 cells infected with short hairpin RNA as a control (Control) or one specific for LOXL2 (LOXL2 KD). Anti-H3K4ox ChIP-PCR was used to analyze the CDH1 promoter in MDA-MB-231 cells infected with either Control (green bar) or LOXL2 KD (blue bar). Data of qPCR amplification were normalized to the input and to total H3 and are expressed as the fold-change relative to data obtained in Control conditions, which were set to 1. (**G**) Western blot of H3K4ox in nucleosomes incubated with recombinant LOXL2 purified from baculovirus, after different incubation times. \*\**P* < 0.01.

### H3K4ox maps to heterochromatin and controls chromatin accessibility in TNBC cells

As aberrant expression and activity of LOXL2 have been reported in various cancers (7-9), we checked the levels of LOXL2 and H3K4ox in several breast cancer cell lines representing different subtypes: luminal A, in the T-47D and MCF-7 cell lines (ER^+^/HER2^−^ /PR^+/–^); luminal B, in the BT-474 cell line (ER^+^/HER2^+^/PR^+/–^); and basal triple-negative breast cancer (TNBC), in the human MDA-MB-231 (ER^−^/HER2^−^/PR^−^) cell line (26). As compared to the other cell lines, MDA-MB-231 (TNBC) showed high levels of LOXL2 and a corresponding enrichment of H3K4ox (Fig. 2A). This trend was observed also in additional TNBC cell lines (e.g., MDA-MB-486, CAL-51, HS-548-T, and BT-549), although with variable LOXL2 expression levels (Fig. 2B). Finally, comparing patient-derived xenographs (PDXs) from luminal (3 PDXs) or TNBC (6 PDXs) subtypes of breast cancer, we found that the levels of both LOXL2 and H3K4ox were higher in the TNBC PDXs (Fig. 2C).

**Fig. 2.**
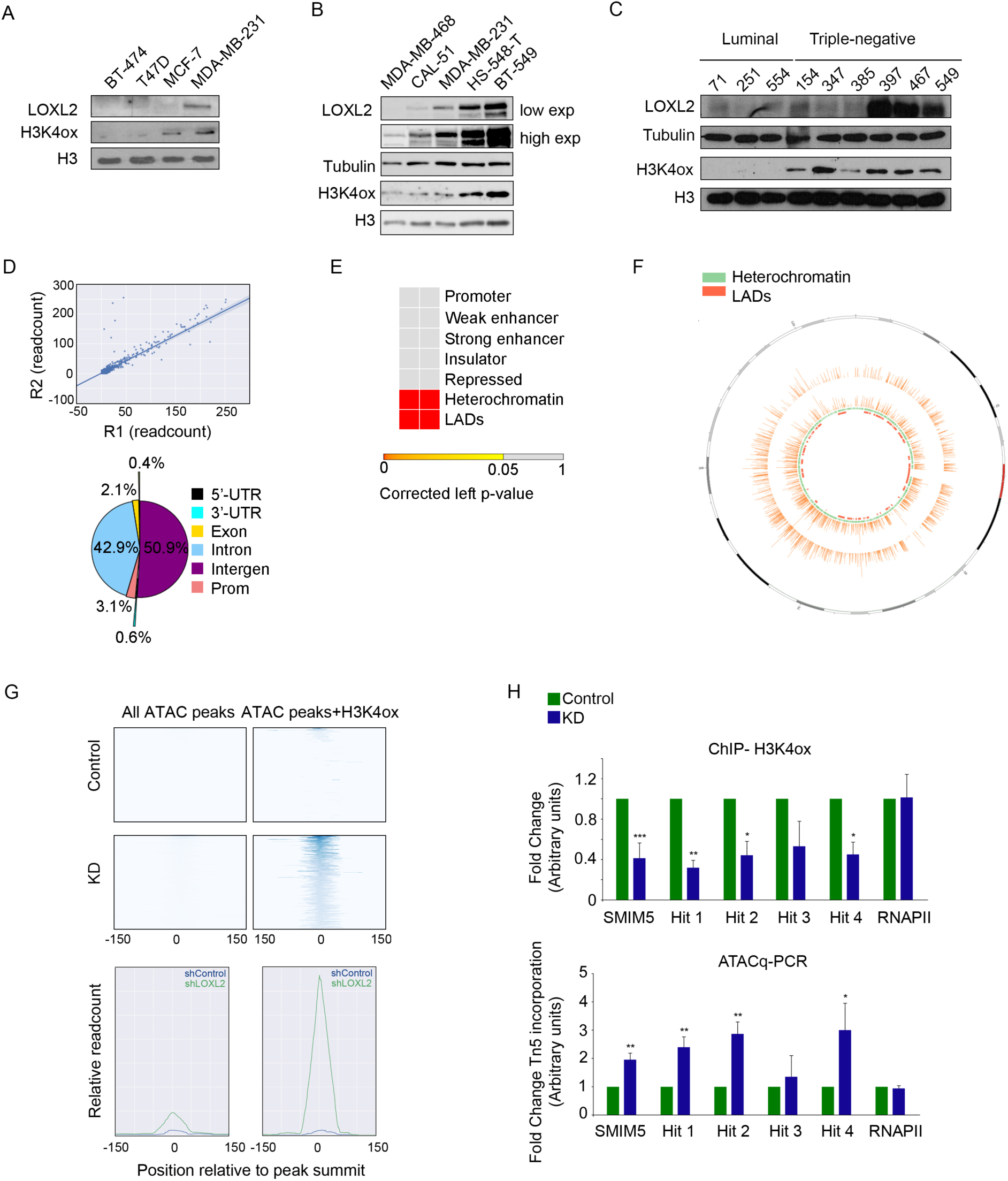
H3K4ox maps heterochromatin and controls chromatin accessibility in TNBC cells. (**A, B, C**) Western blot for the indicated antibodies in a panel of breast cancer cell lines (**A**), TNBC cell lines (**B**), and PDXs (**C**). (**D**) Pearson correlation between two H3K4ox sequencing replicates. Distribution of all H3K4ox ChIP-seq peaks in MDA-MB-231 cells are given, with the indicated percentages. (**E**) Contingency table of the Fisher’s test showed the statistical overrepresentation of the H3K4ox peaks through different chromatin states. (**F**) Circos plot illustrating the location of H3K4ox peaks (orange) in both replicates across chromosome 12, with heterochromatic regions (green) and LADs (red) in the chromosome represented in the innermost tracks. (**G**) Heatmaps show the ATAC signal in all peaks and in peaks that overlap with H3K4ox in LOXL2 KD and control cells. Significant (*P* < 1 × 10^− 5^) ATAC peaks overlapping chromosomic regions (left panel graphs) within significant (*P* < 1 × 10^−5^) H3K4ox ChIP peaks exhibited higher normalized read count than non-overlapping ATAC peaks (right panel graphs). The value of the line in each point is the sum of the read count at the corresponding relative position of the peak, resulting from centering all peaks at their summits. (**H**) H3K4ox ChIP-PCR validation of the selected hits from the ChIP-seq in MDA-MB-231 cells infected with Control or LOXL2 KD. Data of qPCR amplification were normalized to the input and to total H3 and were expressed as the fold-change relative to data obtained in Control conditions, which were set to 1 (upper panel). ATAC-qPCR validation of the incorporation of the transposase Tn5 at the selected hits from the ChIP-seq in MDA-MB-231 cells infected with Control or LOXL2 KD. Data of qPCR amplification were normalized to an unchanging genomic region (HPRT promoter) and expressed as the fold-change relative to data obtained in Control conditions, which were set to 1 (lower panel). RNA polymerase II promoter (RNAPII) was used as a negative control. \**P* < 0.05, \*\**P* < 0.01, \*\*\**P* < 0.001.

To elucidate the function of H3K4ox in breast cancer cells, we first performed a ChIP-seq experiment with the anti-H3K4ox antibody to determine the genomic distribution of H3K4ox in the TNBC cell line MDA-MB-231. Peaks called using model-based analysis for ChIP-seq (MACS) (27) showed low differences in H3K4ox between two sequencing replicates, with a genome-wide Pearson correlation coefficient of the read count of the two replicates of 0.997 (Fig. 2D, upper panel). We observed that H3K4ox peaks were distributed throughout different genomic elements (Fig. 2D, lower panel). Using the chromatin states of two different cell lines determined by the ChromHMM tool (28), we assessed the statistical overrepresentation of the H3K4ox peaks through different chromatin states: promoter, weak or strong enhancer, insulator, repressed, and heterochromatin (non-repetitive sequences) (Fig. 2E). Using the same procedure, H3K4ox peaks were found to be significantly overrepresented in heterochromatin and (as shown previously; *33*) in lamin-associated domains (LADs) in both replicates (Fig. 2, E and F). As generating an aldehyde in H3 removes a positive charge and creates a very reactive group, we hypothesized that this reaction affects chromatin structure. To test this, we used the assay for transposase-accessible chromatin (29,30) followed by deep sequencing (ATAC-seq) that exploits the ability of the prokaryotic transposase Tn5 to integrate preferentially into accessible (open) chromatin. ATAC-seq showed an increased ATAC signal in cells infected with LOXL2 KD, but not with Control, at H3K4ox-positive sites (Fig. 2G). These results were validated in selected regions by ChIP-qPCR and ATAC-qPCR in Control and LOXL2 KD cells: H3K4ox enrichment decreased in LOXL2 knockdown (KD) cells, with a correlating increase of ATAC signal, in these regions (Fig. 2H). No changes were observed in an irrelevant promoter (Fig. 2H). These data demonstrated that, in the absence of LOXL2, H3K4ox levels decrease and chromatin adopts a more open conformation (Fig. 2G). Thus, our results showed that H3K4ox is enriched in heterochromatin and is directly linked with chromatin accessibility in those regions.

### Chromatin structure alterations activate DDR in a LOXL2-dependent manner

As the chromatin state can influence many aspects of the DDR (31), we hypothesized that disruption of LOXL2 expression and impairment of H3K4ox generation influences DDR by affecting chromatin accessibility. To test this, we analyzed MDA-MB-231 cells that had been infected with either LOXL2 KD or Control lentiviruses by immunofluorescence with two well-established markers of DDR: phosphorylated H2AX (γ-H2AX) and TP53-binding protein 1 (53BP1). Depletion of LOXL2 (using LOXL2 KD) led to more foci of both γ-H2AX and 53BP1 than in control cells, suggesting that LOXL2 knockdown (KD) cells may accumulate DNA breaks and/or activate DDR (Fig. 3A). To determine if the catalytic activity of LOXL2 was involved in the observed phenotype, LOXL2 KD-infected cells were complemented by re-infection with ectopic vector expressing either the wild-type (wt) LOXL2-IRES-GFP or a catalytically inactive LOXL2 (LOXL2mut-IRES-GFP), both of which were expressed at similar levels (Fig. S1). Fewer γ-H2AX and 53BP1 foci were observed in LOXL2 KD cells only when wt LOXL2-IRES-GFP was re-introduced (Fig. 3B), establishing that suppressing DDR activation requires both the activity of LOXL2 and H3K4ox generation.

**Fig. 3.**
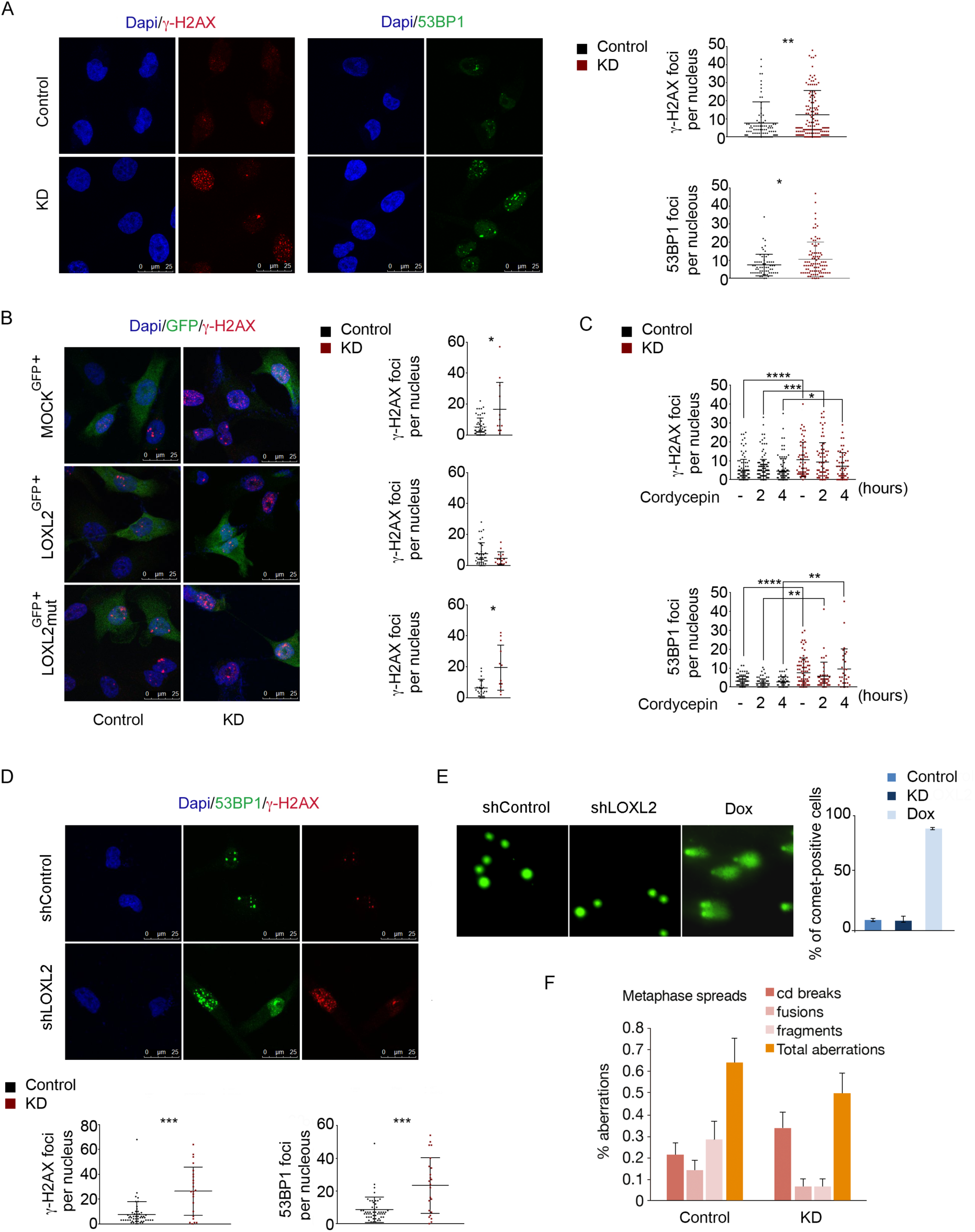
Chromatin opening activates DDR in a LOXL2 catalytic–dependent manner but independently of DNA damage. (**A**) γ-H2AX and 53BP1 staining and foci quantification are shown by immunofluorescence with a specific antibody for γ-H2AX (left image) or for 53BP1 (right image). Dot graphs indicate the number of foci for γ-H2AX (upper graph) and 53BP1 (lower graph) per cell in control and LOXL2 KD conditions. (**B**) γ-H2AX staining and foci quantification are shown by immunofluorescence with the indicated antibody after LOXL2 reinfection. MDA-MB-231 cells were first infected with Control or LOXL2 KD lentivirus and then, after puromycin selection, again with GFP (MOCK^GFP+^), LOXL2-ires-GFP (LOXL2^GFP+^), or LOXL2mut-ires-GFP (LOXL2mut^GFP+^). Cells were fixed after 24 hours. Dot graphs indicate the number of γ-H2AX foci per GFP-positive cell in each condition of MOCK^GFP+^ (upper graph), LOXL2^GFP+^ (middle graph), and LOXL2mut^GFP+^ (lower graph). (**C**) Dot graphs indicate the number of γ-H2AX (upper graph) and 53BP1 (lower graph) foci per cell in Control and LOXL2 KD cells after a treatment with 200 μM cordycepin for the indicated time points. (**D**) γ-H2AX and 53BP1 staining and foci quantification are showed by immunofluorescence with the indicated antibodies in non-replicative conditions. Dot graphs indicate the number of the γ-H2AX (left graph) and 53BP1 (right graph) foci in control and LOXL2 KD cells. **(E**) Representative image showing DNA damage in MDA-MB-231 control or LOXL2 KD cells, visualized by the alkaline comet assay. Cells were treated with 0.3 μM doxorubicin for 24 hours. The graph shows the percentage of MDA-MB-231 positive cells with comet. (**F**) Chromosome alterations in control and LOXL2 KD cells. \**P* < 0.05, \*\**P* < 0.01, and \*\*\**P* < 0.001.

### LOXL2 knockdown activates DDR independently of DNA damage

Increased DDR activation in LOXL2-depleted cells could be a consequence of more DSBs due to reduced H3K4ox levels and/or chromatin decondensation. Notably, aberrant silencing and conflicts between replication forks and transcription, as well as R-loops, a transcriptional intermediate, can result in DNA damage and are influenced by chromatin state, for example in cells lacking the linker histone H1 (32). However, our analysis of RNA-seq data revealed that LOXL2 KD cells did not have altered expression of repetitive elements (Table 1), and DDR activation in LOXL2 KD cells was not affected by cordycepin, an inhibitor of RNA synthesis that abolishes R-loop formation (33,34) (Fig. 3C). As R-loops can generate DNA damage during replication due to fork stalling and collapse (33,35), we next analyzed γ-H2AX and 53BP1 foci in non-cycling cells LOXL2 KD cells and found DDR activation to be normal (Fig. 3D). Overall, these data suggested that overexpression of repetitive elements, R-loop formation, and replication fork stalling were not responsible for activating DDR following LOXL2 depletion, when heterochromatin adopts a more open state. As chromatin structure alterations are responsible to trigger the DDR, even in the absence of DNA damage (19,36,37), we checked for the presence of DNA damage following LOXL2 depletion more directly, using the comet assay, in LOXL2 KD or control cells under alkaline conditions. No increases in DNA damage (due to either single-stranded or double-stranded DNA breaks) were observed in LOXL2 KD cells (Fig. 3E). Moreover, we did not observe any significant differences in chromosomal lesions between LOXL2 KD or control cells in metaphase spreads (Fig. 3F). Finally, analyzing for mitotic aberrations (anaphase bridges and micronuclei) that can be indicative of replication stress or DNA repair defects, we observed only a mild increase in anaphase bridges in the absence of LOXL2 (Fig. S2). Taken together, our data suggested that the combination of loss of LOXL2 and reduced H3K4ox levels in TNBC cells was sufficient to activate the DDR in the absence of detectable DNA lesions or cell cycling defects.

**Table 1.**
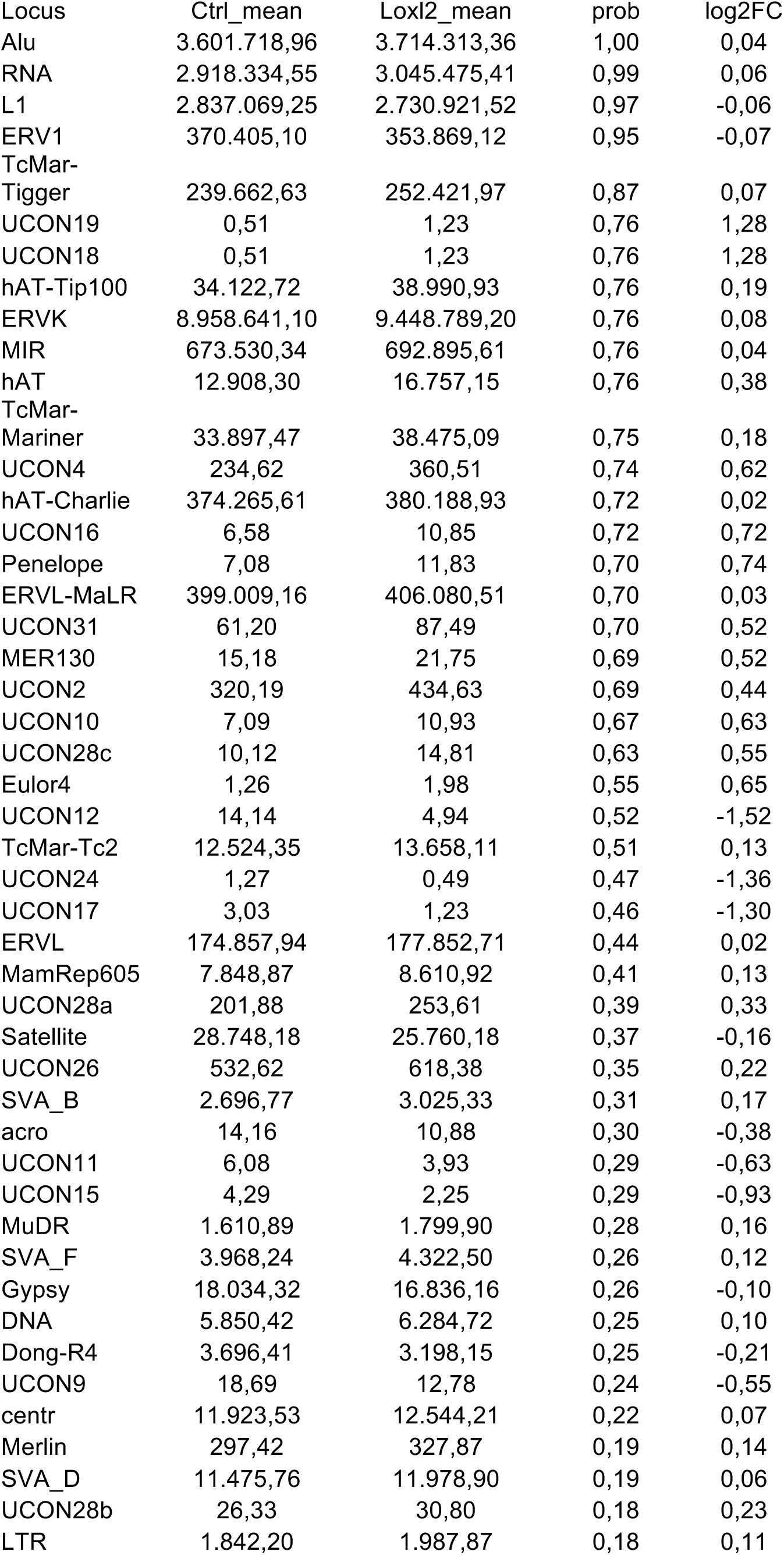
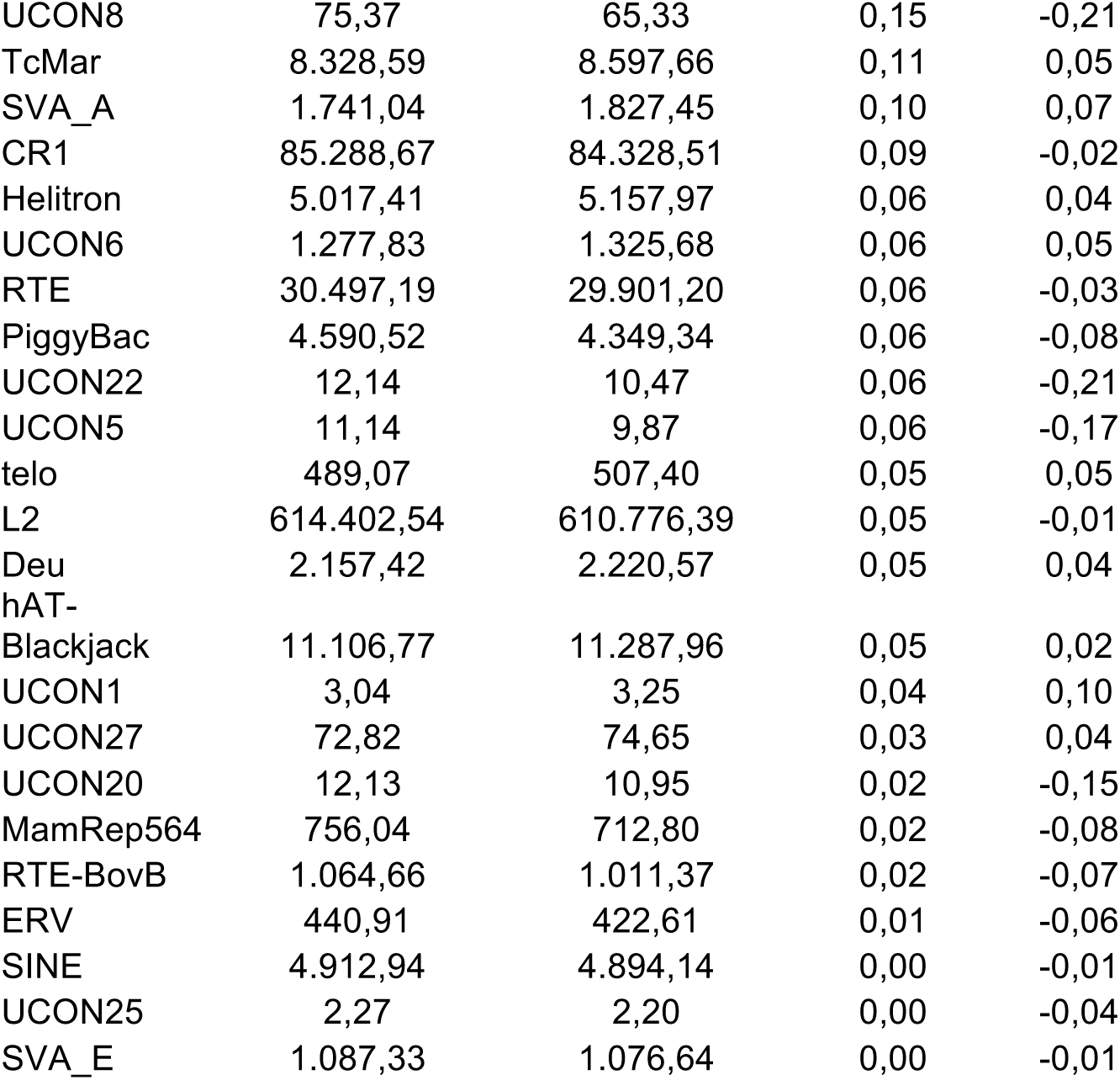
Differential expression analysis of repetitive elements

### Alterations in chromatin compaction activate the DDR

To further address the origin of DDR signaling in LOXL2-depleted cells, we analyzed the behavior of additional DDR signaling components. As both γ-H2AX and 53BP1 foci formation require the ATM kinase in some settings, we treated LOXL2 KD and control cells with the ATM inhibitor KU55933 and analyzed foci formation (Fig. 4A). Decreased foci of both markers were observed upon ATM inhibition, indicating that LOXL2-induced DDR was largely ATM-dependent. Consistent with this, LOXL2 KD cells had increased phosphorylation of several ATM substrates, including KAP-1, CHK1, and CHK2 (Fig. 4B). However, we ruled out that increased DDR signaling was due to apoptosis in LOXL2 KD cells, as no cleaved caspase-3 signal was observed in either LOXL2 KD or control cells (Fig. S3). As these data suggested that the LOXL2 KD cells activated a checkpoint response, we analyzed cell cycle progression following LOXL2 depletion. After synchronization with a double thymidine block, LOXL2 KD cells were not able to efficiently progress through the cell cycle (Fig. 4C), and Western blotting for H3S10-P showed that this histone mark was undetectable in LOXL2 KD cells as compared with control (Control) cells. (Fig. 4D). These data strongly suggested that LOXL2 KD cells arrested primarily in G1; consistent with this possibility, cell proliferation capacity was blocked (Fig. 4E, upper panel) and the colony-formation capacity of LOXL2 KD cells was strongly reduced after only a few passages (Fig. 4E, lower panel).

**Figure 4.**
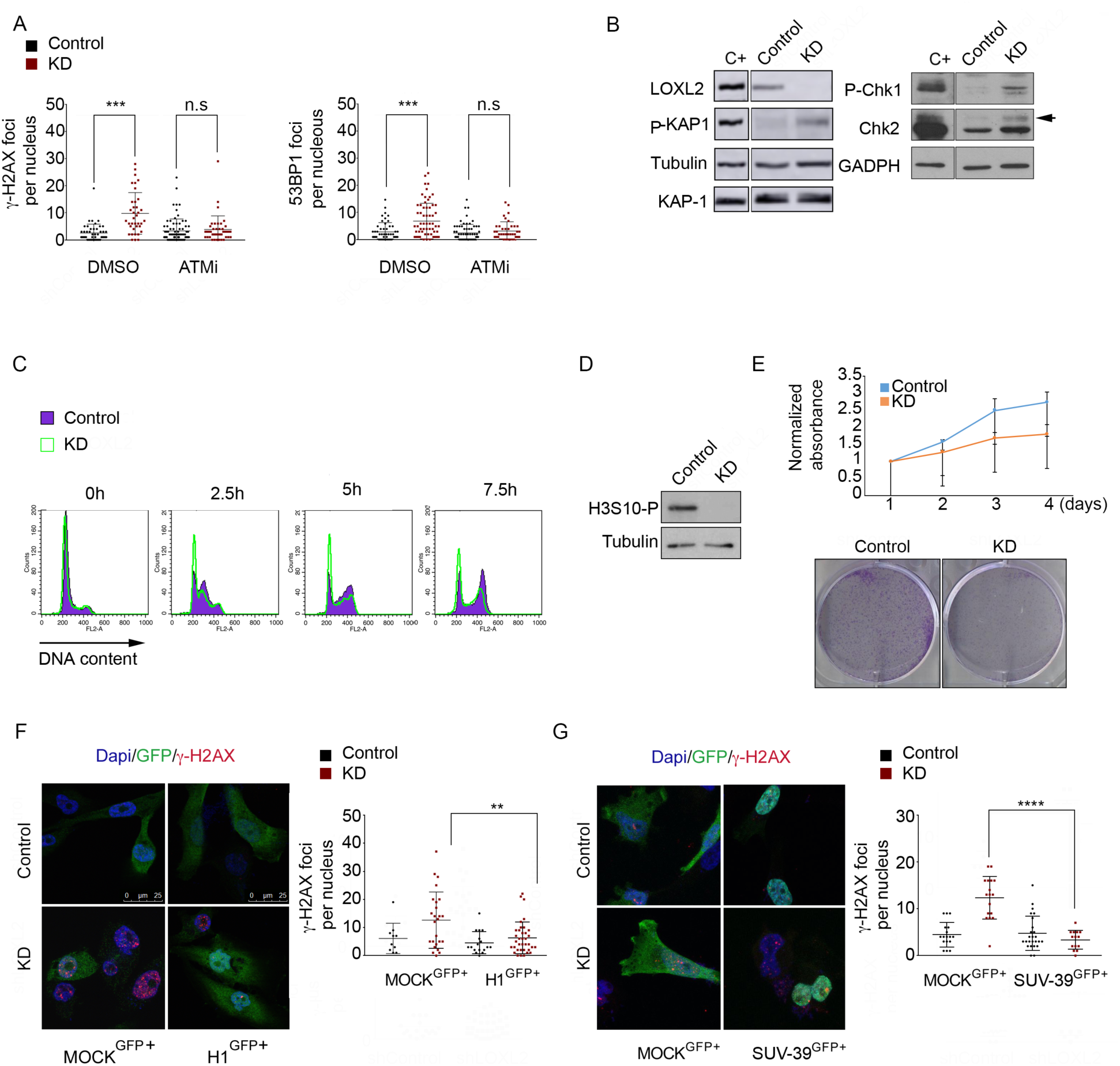
DDR activation is linked to chromatin decondensation in the absence of LOXL2. (**A**) Dot graphs indicate the number of foci with γ-H2AX (left graph) and 53BP1 (right graph) per cell in Control and LOXL2 KD cells treated with DMSO or an ATM inhibitor. (**B**) KAP-1 phosphorylation was analyzed by Western blot in Control and LOXL2 KD cells. Tubulin and total KAP-1 were used as loading controls. As a positive control, MDA-MB-231 cells treated with 0.1 μM doxorubicin for 8 hours were used. Intervening lanes were cropped out as indicated (left panel). Chk1 and Chk2 phosphorylation were analyzed by Western blot in Control and LOXL2 KD cells. Chk1 phosphorylation was detected using the anti-P(S317) Chk1 antibody. For phosphorylated Chk2, a shift was detected using an anti-total Chk2 antibody. GADPH was used as a loading control. All samples were obtained under the same experimental conditions; in addition, positive control samples (irradiated fibroblasts) were run on the same gel as their corresponding Control and LOXL2 KD samples. Intervening lanes in the Chk1/2 and GADPH were cropped out as indicated (right panel). (**C**) Cell cycle profile of Control and LOXL2 KD cells at 0, 2.5, 5, or 7.5 hours after release from a double-thymidine block. Cells were analyzed by FACS after propidium iodide staining. (**D**) H3S10 phosphorylation levels were analyzed by Western blot in Control and LOXL2 KD cells MDA-MB-231 cells. Tubulin was used as a loading control. (**E**) Upper panel, MTT assay in in Control and LOXL2 KD MDA-MB-231 cells; Lower panel, Colony-survival assay in Control and LOXL2 KD MDA-MB-231 cells. (**F** and **G**) γ-H2AX staining and foci quantification are shown by immunofluorescence with the indicated antibody after H1 reinfection (**F**) or SUV-39H1 reinfection (**G**). MDA-MB-231 cells were infected with Control or LOXL2 KD lentivirus, selected with puromycin, and re-infected with either GFP (MOCK^GFP+^) or with histone 1-GFP (H1^GFP+^) (**F**) or SUV-39H1-GFP (SUV-39^GFP+^) (**G**). Cells were fixed after 24 hours. Dot graphs indicate the number of γ-H2AX foci per GFP-positive cell in each condition. \**P* < 0.05, \*\**P* < 0.01, \*\*\**P* < 0.001.

To test whether the effects on chromatin in LOXL2 KD cells directly activated ATM-dependent DDR signaling, we forced chromatin condensation in these cells by expressing the linker histone H1 or the H3K9 methyltransferase SUV-39H1 (as GFP-labeled proteins) and counted the number of γ-H2AX foci in GFP positive cells. Notably, overexpression of either H1^GFP^ or SUV-39 ^GFP^ reduced the number of γ-H2AX foci in LOXL2 KD cells as compared to LOXL2 KD cells that expressed GFP alone (+MOCK^GFP^) (Fig. 4F and G). This suggested that lack of LOXL2 and reduced H3K4ox levels affected the regulation of chromatin condensation (leading to decondensation) and activated DDR, even in the absence of DNA damage.

### LOXL2 reduction enhances chemosensitivity of TNBC cells

As we found that reducing H3K4ox levels led to chromatin decondensation, triggered DDR activation, and induced cell cycle arrest, we wondered whether having higher LOXL2 levels would reduce the cellular response to DNA-damaging chemotherapy (thereby making cancer cells less sensitive to chemotherapy). However, as no LOXL2 inhibitors are currently available, we first tested whether TNBC cell lines with low, medium, or high LOXL2 expression levels responded distinctly to treatment with doxorubicin, a topoisomerase inhibitor that generates DSBs. Indeed, we observed that a LOXL2 high-expression cell line, BT-549, was more resistant to doxorubicin than the MDA-MB-231 and MDA-MB-468 cell lines (which have lower LOXL2 levels). However, in all cases, treatment of doxorubicin together with trichostatin A (TSA), a general HDCA inhibitor (38) that increases chromatin accessibility (39), increased the percentage of cell death—indicative of an increased cell sensitivity to chemotherapy (Fig. 5A). Similar results were obtained with two TNBC_PDXs: PDX-549, which has higher levels of LOXL2, was more resistant to chemotherapy than PDX-154 (Fig. 5A). To determine if these results could be reproduced in vivo, PDX-549 cells were first grown ex vivo and then 10^6^ cells were subcutaneously implanted in Nude mice. After tumor formation, mice were treated with TSA, doxorubicin, doxorubicin plus TSA, or (as a control) vehicle for 25 days. In agreement with our previous results, we observed that tumor growth was substantially impaired with the combined TSA/doxorubicin treatment (Fig. 5B).

**Figure 5.**
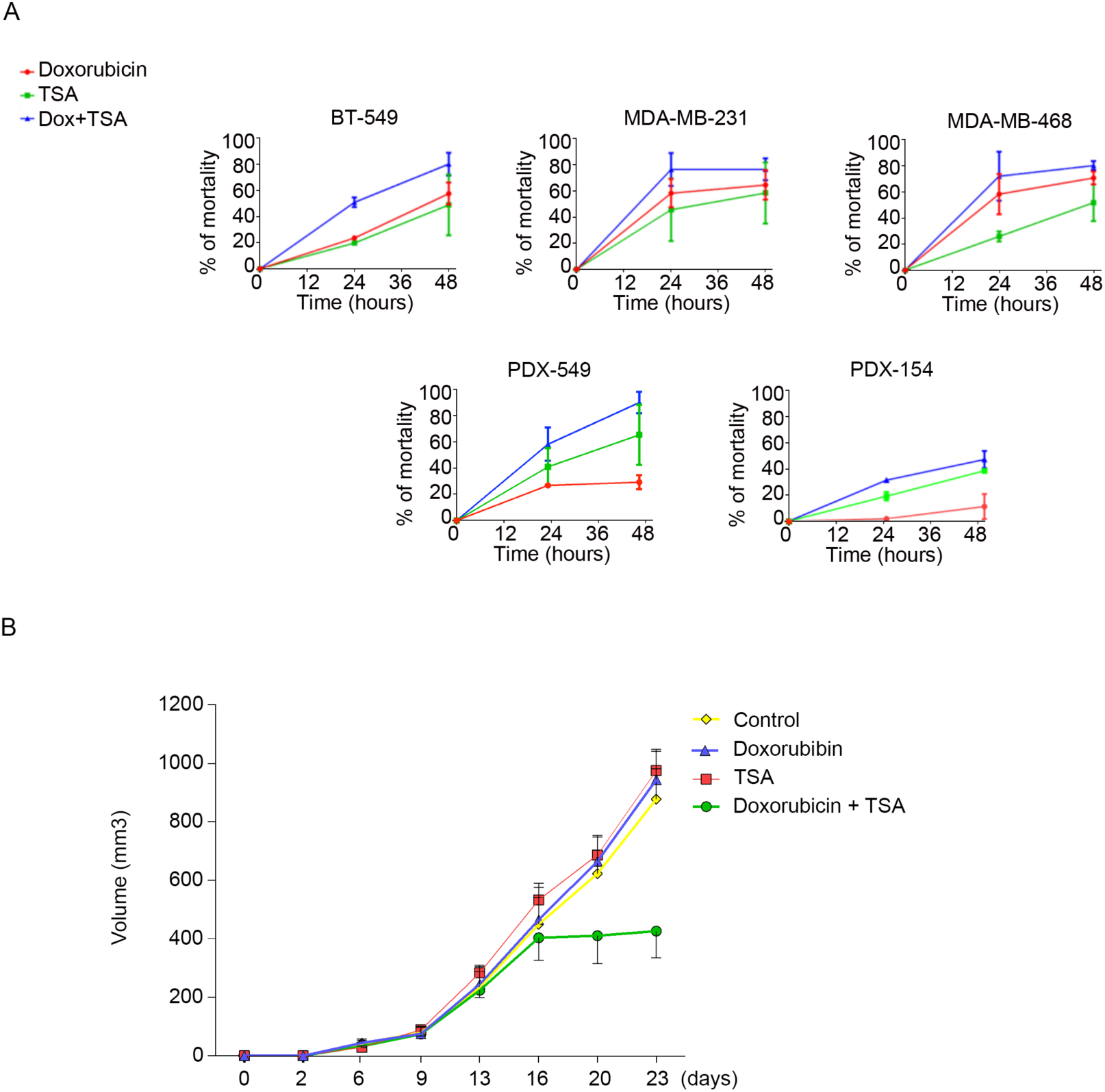
LOXL2 suppresses sensitivity in TNBC cells. (**A**) Cell viability was measured at different time-points using an MTT assay of cultured different breast cancer cells and PDXs. The effects of doxorubicin, TSA, and doxorubicin plus TSA were analyzed. (**B**) Dissociated cells from PDX-549 were orthotopically implanted into NOD/SCID mice and injected intraperitoneally twice weekly with TSA (0.25 mg/kg), doxorubicin (2 mg/kg mouse weight), or doxorubicin plus TSA. Tumor volumes were measured twice a week and are given as averages. Results are given as averages of six independent tumors ± SEM. \**P* < 0.05, \*\**P* < 0.01, \*\*\**P* < 0.001

## Discussion

In this work, we found that many TNBC cell lines and PDXs express high levels of LOXL2 as compared to the luminal breast cancer subtype. With the use of a newly-generated antibody, we were able to detect the H3 modification produced by LOXL2, H3K4ox (3). This H3 modification is enriched in heterochromatin, where it is required to maintain a condensed state. It is possible that during tumor progression, some cancer cells undergo the EMT program and start to express LOXL2. The transcription factor Snail1, together with LOXL2, would participate in downregulating the CDH1 gene and heterochromatin transcripts, giving rise to transformation of cancer epithelial cells into mesenchymal cells (through EMT) (6,40-42). Recently, two different groups have suggested that, although EMT is dispensable for lung and pancreas metastasis, it contributes significantly to the formation of recurrent metastasis after chemotherapy (43,44). This finding is in agreement with our results, in which we observed that the most aggressive and resistant subtype of breast cancer (TNBC) have a mesenchymal phenotype and enriched H3K4ox levels in heterochromatin. We observed that reducing H3K4ox levels perturbed the balance between condensed and decondensed chromatin and activated the DDR. Similar phenotypes have been previously observed after depletion of SUV-39, PRDM2, or HP1, which are required for heterochromatin maintenance and cell survival after DNA damage (20,22,45,46). Moreover, sustained upstream DDR signaling was observed when chromatin is maintained in a condensed state (19,47,48). We found that forcing chromatin decondensation activated DDR signaling in the absence of detectable DNA lesions. As LOXL2 has multiple substrates besides histones (5), we cannot discard an additional putative role for LOXL2 in the direct regulation of the DDR machinery. However, we observed that DDR induced in the absence of LOXL2 can be inhibited by forcing the compaction of chromatin, suggesting that the main molecular mechanism is the regulation *per se* of the chromatin compaction state.

While a correlation between chromatin compaction and DDR has been previously reported (19,47,48), we demonstrated here that H3 oxidation by LOXL2 is another molecular mechanism that maintain compaction. Notably, LOXL2 has been widely linked to cancer and the acquisition of cellular malignancy, as it is overexpressed in many tumors (15,49-52) and has an important role in tumor formation (53). Our analyses of LOXL2 levels in cell lines and PDXs, revealed that LOXL2 and H3K4ox are mainly expressed in the TNBC subtype of breast cancer. These results suggest that LOXL2/H3K4ox levels could be an indicator of patient response to chemotherapy.

Induction of chromatin compaction has been suggested as a potential therapeutic tool in gliomas, in which chromatin compaction limits DDR (47) and promotes damage resistance (54). However, this does not appear to apply to TNBC cells. In support of this, we have now shown that several TNBC PDX tumors express higher levels of LOXL2 and higher H3K4ox levels relative to other TNBC PDXs, and that treatment with TSA, a histone deacetylase inhibitor that generates an open chromatin state, enhanced tumor killing with doxorubicin in these cells. These observations, together with previously published clinical data, suggest a rationale for studying combinations of “chromatin-opening drugs”, including inhibitors of histones deacetylases, EZH2, or LSD1, in TNBC to overcome resistance to treatment.

## Materials and Methods

### Cell lines, transfections, and infections

Most cell lines (the TNBC lines MDA-MB-468, CAL-51, HS-548T, and MDA-MB-231, the luminal A T-47D, MCF-7, and luminal B BT-474) as well as cells isolated from PDXs were maintained in Dulbecco’s modified Eagle’s medium (Biowest; L0106-500), and the TNBC line BT-549 was maintained in Roswell Park Memorial Institute medium (Biowest; L0501-500); all were supplemented with 10% fetal bovine serum (Gibco; 10270106), 2 mM L-glutamine (Biowest; X0550-100), and 1% penicillin-streptomycin (Gibco; 15140122) at 37°C in 5% CO_2_. For LOXL2-Flag overexpression assays, MCF-7 cells were seeded for 24 hours and transfected with 10 μg pcDNA3-hLOXL2-Flag vector using polyethylenimine polymer (Polysciences Inc; 23966-1). For SUV39H1-GFP rescue experiments, MDA-MB-231 cells under selection were seeded for 24 hours and transfected with 2.5 μg CGA-pCAGGS-Suv39h1-EGFP-IRES-Puro vector using TransIT-X2 Dynamic Delivery System (Mirus Bio LLC; MIR600Q). For lentiviral infections to produce LOXL2 KD cells, HEK293T cells were used to produce lentiviral particles. Cells were grown to 70% confluency and then transfected by dropwise addition of NaCl, DNA composed of 50% pLKO-shCT/LOXL2 KD, 10% pCMV-VSVG, 30% pMDLg/pRRE, 10% pRSV rev, and polyethylenimine polymer (Polysciences Inc), which had been pre-incubated for 15 min at room temperature; transfection medium was replaced with fresh medium after 24 hours. Viral particles were concentrated using Lenti-X Concentrator product (Clontech; 631232) and then used to infect MDA-MB-231 cells (Millanes-Romero et al., 2013). For retroviral infections, HEK293 gag-pol cells were used to produce retroviral particles. Cells were transfected as for HEK293T cells with a DNA mixture of DNA (2.5 μg of pCMV-VSV-G and 7.5 μg of pMSCV, pMSCV-LOXL2 wt-FLAG or pMSCV-LOXL2 mutFLAG IRES-GFP vectors) and polyethylenimine polymer (Polysciences Inc; 23966-1) that had been pre-incubated for 15 min at room temperature; transfection medium was replaced with fresh medium after 24 hours. Viral particles were concentrated using Retro-X Concentrator product (Clontech; 631456) and then used to infect MDA-MB-231 cells. HEK293T cells were used to produce lentiviral particles for H1-GFP expression. Cells were first grown to 70% confluency and transfected with a mixture of NaCl, DNA composed of 50% FUGW-H1-empty vector/FUGW-H1-GFP, 10% pCMV-VSVG, 30% pMDLg/pRRE, and 10% pRSV rev, and polyethylenimine polymer (Polysciences Inc; 23966-1), which had been pre-incubated for 15 min at room temperature; transfection medium was then replaced with fresh medium after 24 hours. Viral particles were concentrated using Lenti-X Concentrator product (Clontech; 631232) and used to infect MDA-MB-231 cells.

### Antibodies and other reagents

The following antibodies were used: anti-FLAG (F7425, Sigma), anti-LOXL2 (NP1-32954, Novus), anti-H3K4me3 (07-473, EMD Millipore), anti-H3K9me3 (07-442, Millipore), anti-phospho-histone H2AX (S139) clone JBW301 (05-636, EMD Millipore) and for ChIP (ab2893, Abcam), anti-GFP (ab6556, Abcam), anti-53BP1 (NB100-904, Novus Biol.) and for ChIP (NB100-304, Novus Biol.), anti-phospho-CHK1 (S317) (A300-163A, Bethyl), anti-CHK2 clone 7 (05-649, EMD Millipore), anti-cleaved caspase 3 (Asp175) (9661, Cell Signaling), anti-KAP1 (ab10484, Abcam), anti-Phospho-KAP1 (S824) (A300-767A, Bethyl), and anti-histone H3 (ab1791, Abcam). The following chemical reagents were used in several experimental conditions: doxorubicin hydrochloride (Sigma; D1515), cordycepin (Sigma; C3394) and the ATM inhibitor KU55933 (Sigma; SML1109). An anti-H3K4ox antibody was generated from a synthetic peptide in which Fmoc-6-hydroxynorleucine (BAA1117, Iris Biotech) instead of Fmoc-lysine(Boc) was incorporated at position 4. The peptide was purified (>95%) by RP-HPLC, its identity confirmed by MS, then it was coupled to keyhole limpet hemocyanin (KLH) for antibody production.

### Cell extracts, histone isolation, and PDX extraction

To obtain nuclear fractions of LOXL2-Flag–transfected MCF-7 cells and HEK293T cells, cells were first lysed in soft-lysis buffer (50 mM Tris, 2 mM EDTA, 0.1% NP-40, and 10% glycerol, supplemented with protease and phosphatase inhibitors) for 5 min on ice. Samples were centrifuged at 3,000 rpm for 15 min, and the supernatant was discarded. The nuclear pellet was lysed in high-salt lysis buffer (20 mM HEPES pH 7.4, 350 mM NaCl, 1 mM MgCl_2_, 0.5% Triton X-100, and 10% glycerol, supplemented with protease and phosphatase inhibitors) for 30 min at 4°C, and samples were centrifuged at 13,000 rpm for 10 min. Supernatant NaCl concentration was reduced to 300 mM NaCl by adding balanced buffer (20 mM HEPES pH 7.4, 1 mM MgCl_2_, and 10 mM KCl). Cells extracts from the different breast cancer cell lines were obtained with SDS lysis buffer (2% SDS, 50 mM Tris-HCl, and 10% glycerol). Histones were isolated from the different breast cancer cell lines using acid precipitation. For this, cell monolayers were first scraped in 1 ml of lysis buffer (10 mM Tris pH 6.5, 50 mM sodium bisulfite, 10 mM MgCl_2_, 1% Triton X-100, 8.6% sucrose, and 10 mM sodium butyrate). Working at 4°C, pellets were then purified by three rounds of centrifugation (13,000 rpm for 15 sec) and resuspension by vortexing, in lysis buffer for the first two rounds and washing buffer (10 mM Tris pH 7.4, 13 mM EDTA) for the final round; supernatant fractions were discarded. The obtained pellets containing chromatin were resuspended in 0.4 N sulfuric acid, the mixture was incubated at 4°C for 1 hour and then centrifuged at 13,000 rpm for 10 min, and the histone-containing supernatants were kept. Samples were incubated with acetone (1:9) to block the acid overnight at –20°C and then centrifuged at 13,000 rpm for 10 min. The histone pellets were air dried for 5 min (supernatant fractions were discarded). Pellets were resuspended in water for analysis. Cell extracts of PDXs samples were obtained by disrupting tissue using a Pellet Pestle (Sigma; Z359947) with lysis buffer (50 mM Tris-HCl pH 8, 10 mM EDTA, 1% SDS, and 1 mM DTT). Proteins were separated by SDS-polyacrylamide gel electrophoresis gel and analyzed with the indicated antibodies.

### Recombinant LOXL2 purification

LOXL2-encoding baculovirus were amplified, and LOXL2-Flag recombinant proteins (wild-type and mutant) proteins were produced in Sf9 cells according to standard procedures (3). Cell lysis was performed as previously described (55). Cell extracts were incubated with Flag M2 beads for 4 hours at 4°C and then washed 4× times with 20 mM HEPES pH 7.4, 1 mM MgCl_2_, 300 mM NaCl, 10 mM KCl, 10% glycerol, and 0.2% Triton X-100. Recombinant proteins were eluted with the Flag peptide (1 μg/μl) for 1 hour at 4°C.

### Dot blot assay

For dot blot assays, 1 μg of each peptide (in 10 μl of sample) was applied under low vacuum to a pre-wetted nitrocellulose membrane (Amersham Protran 0.45 nitrocellulose, GE Healthcare) using a dot blot apparatus (HYBRI-DOT Manifold; Life Technologies). The entire blot was blocked in 15 mL of 5% non-fat dry milk and 0.1% Tween-20 Tris-buffered saline for 1 hour at room temperature and then probed with the indicated antibodies.

### ChIP experiments

For ChIP experiments, cells were first crosslinked in 1% formaldehyde for 10 min at 37°C. Crosslinking was stopped by adding glycine to a final concentration of 0.125 M for 2 min at room temperature. Cell monolayers were scraped in cold soft-lysis buffer (50 mM Tris pH 8.0, 10 mM EDTA, 0.1% NP-40, and 10% glycerol), and incubated 20 min on ice. Nuclei pellets were lysed with SDS lysis buffer (1% SDS, 10 mM EDTA, 50 mM Tris pH 8.0), and extracts were sonicated to generate 200- to 1,500-bp DNA fragments. For immunoprecipitation, supernatants were diluted 1:10 with dilution buffer, and samples were incubated with rotation overnight at 4°C with primary antibody or irrelevant IgGs. Samples were then treated with elution buffer (100 mM Na2CO3 and 1% SDS) for 1 hour at 37 °C and incubated at 65°C overnight after addition of NaCl to a final concentration of 200 mM, to reverse formaldehyde crosslinking. After proteinase K treatment for 1 hour at 55°C, DNA was purified with MinElute PCR purification kit (Qiagen; 28006) and eluted in Milli-Q water. Genomic regions were detected by quantitative PCR SYBR Green staining (Quantabio; 95073), and the ChIP results were quantified relative to the input amount and the amount of H3 immunoprecipitated in each condition.

Peaks of H3K4ox were called from sequence reads detected through ChIP-seq using the MACS2 tool (27). The chromatin state files for Hpeg2 and HMEC cells were computed by the ENCODE project using the ChromHMM tool (28) from https://genome.ucsc.edu/cgi-bin/hgFileUi?db=hg19&g=wgEncodeBroadHmm. Statistical overrepresentation of H3K4ox peaks detected by ChIP-seq was assessed across several chromatin states from these two cell lines: heterochromatin, repressed, insulator, strong enhancer (sum of states 4 and 5), weak enhancer (sum of states 6 and 7), promoter (sum of states 1 and 2), and poised promoter. The contingency table of the Fisher’s test carried out to this end contained the number of nucleotides within peaks, chromatin states, intersections thereof, and the remaining portion of the genome (computed as the difference from the effective genome size for ChIP-seq peaks calling). The same procedure was applied to detect H3K4ox peak overrepresentation in lamin-associated domains of chromatin, obtained from (56).

### ATAC-seq and ATAC-qPCR experiments

ATAC experiments were performed as described previously (29). Cells were harvested and treated with transposase Tn5 (Nextera DNA Library Preparation Kit, Illumina; FC-121-1030). DNA was purified using MinElute PCR purification kit (Qiagen), samples were amplified by PCR using NEBNext High-Fidelity 2× PCR Master Mix (New England Biolabs; M5041), and DNA was again purified with the MinElute PCR purification kit. Finally, qPCR was performed with the same primers as for the ChIP experiments on a 1/50 dilution of the eluted DNA. Reads produced by ATAC sequencing of two control (Control) replicates and two LOXL2 knockdown sequencing replicates (LOXL2 KD) were aligned to the hg19 build of the reference human genome using Bowtie 2 (57) with default parameters for pair-end sequencing. ATAC peaks were then called by combining aligned reads of both replicates of the control and the knockdown using MACS2. To allow for FDR threshold selection further downstream in the analysis, no FDR restrictions were imposed on the ATAC peak calling.

### RNA-seq analysis

Reads produced by RNA-seq of the same two control (Control) and two LOXL2 knockdown (LOXL2 KD) replicates as above were aligned to the hg19 build of the reference human transcriptome using TopHat2 (58) with default parameters for pair-end sequencing. Aligned reads were then analyzed using a standard Cufflinks pipeline (59) to detected differentially expressed genes between the two conditions (LOXL2 KD and Control).

### Differential expression analysis of transposable elements (TEs)

The trimmed RNA-seq reads from the two controls and the two LOXL2 samples were processes with the TEtools program (60) using the hg19 annotation of TEs. The obtained count table was then processed with NOIseq (61) to perform a differential expression analysis of LOXL2 against the control. Expression was normalized using the TMM method, whereby TEs with a probability higher than 0.95 of being differentially expressed were considered as statistically significant.

### Replicate correlation

The read count (coverage) at each position of the hg19 human reference genome was computed for each replicate of the H3K4ox ChIP-seq and the Control and LOXL2 KD ATAC sequencing, using the BEDtools *genomecov* capability (62). Genomic positions with zero read counts were filtered out. Replicate files of each experiment were merged to produce a single file aligned by genomic position, and the corresponding Pearson’s correlation coefficient of read counts was computed. For the graphical representation of the correlation, 100,000 genomic positions were randomly selected.

### Analysis of ATAC peaks that overlap with H3K4ox peaks

All significant (*P* < 10^−5^) ATAC peaks (LOXL2 KD versus Control) and H3K4ox peaks were first intersected with the BEDtools *intersect* program (62). Based on this intersection, ATAC peaks were classified as *overlapping* (if they intersected an H3K4ox peak) or *orphan* (if not). Only intersections involving more than 95% of the sequence of ATAC peaks were considered. Control and LOXL2 KD read counts over all genomic positions (see above) were intersected with both overlapping and orphan peaks. Read counts over genomic positions of control and experiment replicates were averaged. To carry out the heatmap representation, peak sequences (overlapping or orphan) were aligned by their summits. For linear representation, the average experimental read counts at each downstream and upstream position were summed for both the experimental and the control counts. Position-wise sums were then divided by the read count sum value obtained for the summit of control read counts, thus making all sums relative to the maximum control value.

### Integrated analysis of H3K4ox and ATAC Peaks and Differentially Expressed Genes

The differentially expressed genes detected through the RNA-seq analysis of Control and LOXL2 KD cells were selected if they were close (up- or downstream) to *overlapping* ATAC peaks. Two different distance thresholds were used to detect close differentially expressed genes: 0.5 Mb and 1 Mb.

### Cordycepin inhibition of RNA synthesis

MDA-MB-231–infected cells under selection were seeded for 48 hours after selection. Cells were then incubated with 200 μM cordycepin (Sigma; C3394) and fixed at designed time points.

### KU55933 inhibition of ATM kinase

MDA-MB-231–infected cells under selection were seeded for 24 hours after selection. Cells were incubated for 24 hours in two different conditions 5 μM and 10 μM KU55933 (Sigma; SML1109). As a positive control, 1 μM doxorubicin was added for 2 hours.

### Cell cycle analysis

MDA-MB-231–infected cells under selection were first synchronized through the double thymidine block protocol. Cells were seeded to 50% confluency, incubated for 14 hours with complete growth medium supplemented with 2 mM thymidine, washed 2× with PBS, and released by a 9-hour incubation with complete medium growth. Cells were washed again 2× with PBS, incubated 14-hour with 2 mM thymidine, and released with complete growth medium. Cells were then harvested at the designed time points and fixed with 100% cold ethanol. After two days, fixed cells were stained with propidium iodide (PI) staining and analyzed by flow cytometry using BD FACSCalibur (Becton Dickinson). Results were analyzed using BD CellQuest Pro software.

### Non-replicative cell experiment

MDA-MB-231 cells were seeded in coverslips and maintained during all the experiment in Dulbecco’s modified Eagle’s medium (Biowest; L0106-500) with 0.5% fetal bovine serum (Gibco; 10270106) at 37°C in 5% CO_2_. After 24 hours, cells were infected with lentiviral particles for LOXL2 knock down. After 96 hours under selection, cells were fixed.

### Comet assay

MDA-MB-231–infected cells under selection were seeded for 48 hours after selection.

### Rescue experiments

MDA-MB-231–infected cells under selection were seeded for 24 hours after selection. Cells were then transfected with the SUV39H1-EGFP vector or re-infected with retroviral particles for LOXL2-FLAG or lentiviral particles for H1 expression. After 24 hours, cells were fixed for immunofluorescence.

### Immunofluorescence, image acquisition, and analysis

Cells were fixed with 4% PFA for 15 min at room temperature, blocked for 1 hour with 1% PBS-BSA, incubated at room temperature for 2 hours with primary antibody, washed 3× with PBS, and then incubated for 1 hour at RT with the secondary antibody. Cells were washed again 3× with PBS, incubated for 5 min with DAPI (0.25 mg/ml) for cell nuclei staining, and then mounted with fluoromount. Fluorescence images corresponding to DAPI, γ-H2AX, 53BP1, and H3K4ox were acquired in a Leica TCS SPE microscope using a Leica DFC300 FX camera and the Leica IM50 software.

### Metaphase spreads

For metaphase spread preparations, cells were treated with colcemid (0.1 g/ml for 4 h). Cells were trypsinized, hypotonically swollen in 0.075 M KCl for 15 min at 37 °C, and then fixed (75% MeOH and 25% acetic acid, ice cold). Metaphase preparations were spread on glass slides, stained with 10% Giemsa stain (Sigma), and mounted in DPX mounting medium (PanReac). Images were taken using a Leica DM6000 microscope (Leica, Wetzlar, Germany) and analyzed using Fiji Software (https://fiji.sc/).

### Cellular viability experiment

Breast cancer cell lines and PDX cells were seeded in 96-well plates. Cellular viability was analyzed using Thiazolyl Blue Tetrazolium Bromide (MTT) (Sigma; M5655) at different time points. Concentrations of 0.1 μM doxorubicin for all samples, 250 nM TSA for the breast cancer cell lines, and 500 nM TSA for the PDXs cells were used. Absorbance was detected at 565 nm on Infinite^®^ 200 PRO Series Multimode Reader (Tecan Group Ltd.) and analyzed with i-control(tm) Microplate Reader Software (Tecan Group Ltd.)

### Cancer patient-derived xenografts (PDXs) and treatments

PDXs were obtained from the operation room and transferred to the pathology department, where breast cancer samples were collected, transferred to the animal facility, and implanted into mice. All samples were implanted within 60 to 90 min after surgical removal. All patients signed an informed consent, and the study was approved by the Ethics Committee of the Vall d’Hebron Hospital.

For in vivo experiments, a tumor from an established PDX was dissociated into single cells by enzymatic digestion (collagenase at 300U/ml and hyaluronidase at 200U/ml) during 1 hour at 37°C in a rotating wheel. The solution of digested tumor was treated with 0.025% trypsin and filtered sequentially using 100 μm and 40 μm strainers. Isolated cells were plated in culture dishes with DMEM-F12 supplemented with FBS, glutamine, and penicillin/streptomycin. Once the culture was established, 10^6^ cells were injected into the number four fat pad of six-week old NOD.CB17PrkdcSCID/J (NOD/SCID) females (Charles River) with Matrigel. For this, mice were anesthetized and shaved, the fourth and fifth sets of nipples were localized, and an inverted Y incision was made from the midline point between the fourth set of nipples, ending between the fourth and fifth sets to expose the fourth and fifth fat pads on one side. After the injection, animals were sutured, and analgesics injected. Animals were kept in a clean cage with drinking water supplemented with 1 μM 17-β-estradiol. Tumor xenografts were measured with callipers every 3 days, and tumor volume was determined using the formula: (length × width^2^) × (pi/6). At the end of the experiment, animals were anesthetized with a 1.5 % isofluorane-air mixture and were sacrificed by cervical dislocation.

Treatments were administered intraperitoneally twice weekly. 1 week after cell injection, mice were randomized and treated with TSA (0.25 mg/kg), doxorubicin (2 mg/kg mouse weight), or doxorubicin plus TSA. The control group was injected with sterile PBS.

Mice were maintained and treated in accordance with institutional guidelines of Vall d’Hebron University Hospital Care and Use Committee.

## General

Thank R. Peña and J. Valle for technical assistance, V. Raker for manuscript editing, G. Gil for manuscript reading and advice, A. Jordan for H1-GFP constructs, T. Jenuwein for SUV-39 constructs and H. Galvez-Garcia for ATAC protocol implementation, thers for any contributions.

## Author contributions

Describe the contributions of each author (use initials) to the paper.

## Data and materials availability

GSE96064

## Supplementary Materials

**Supplementary Figure 1.**
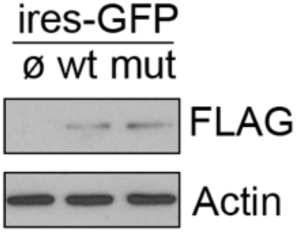
Western blot with the indicated antibodies in 293 cells infected with GFP (MOCKGFP+) or with LOXL2-IRES-GFP (LOXL2GFP+) or LOXL2mut-IRES-GFP (LOXL2mutGFP+).

**Supplementary Figure 2.**
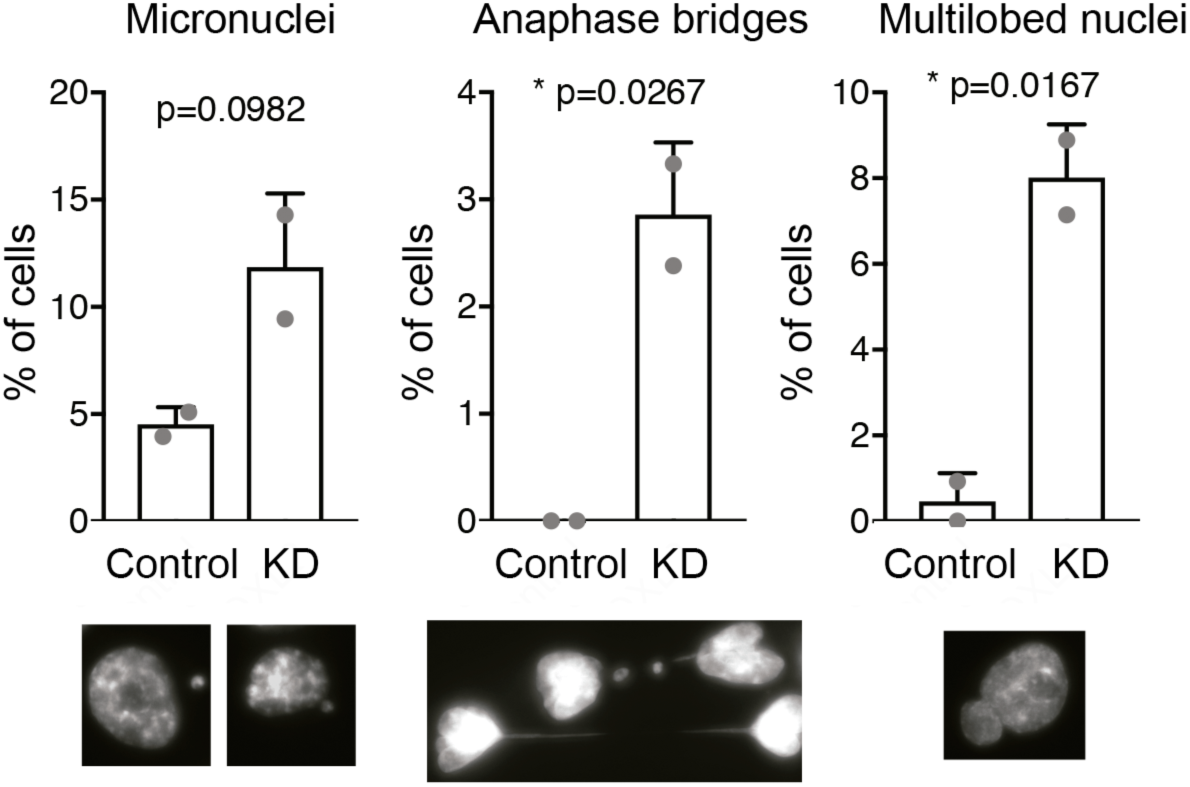
Percentage of the indicated chromosome alterations per total number of cells in shControl and shLOXL2 conditions in two independent experiments. *p<0.05, **p<0.01, ***p<0.001.

**Supplementary Figure 3.**
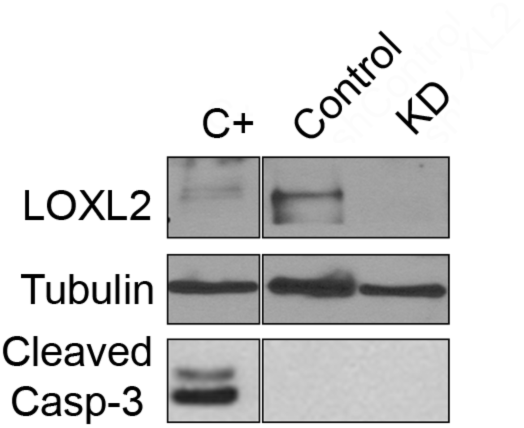
Caspase-3 activation analyzed by Western blot with the indicated antibodies. Tubulin was used as a loading control. Intervening lanes were cropped out as indicated.

## References and Notes

1. Bannister, A.J. and Kouzarides, T. (2011) Regulation of chromatin by histone modifications. Cell research, 21, 381–395.

2. Williamson, P.R. and Kagan, H.M. (1986) Reaction pathway of bovine aortic lysyl oxidase. J Biol Chem, 261, 9477–9482.

3. Herranz, N., Dave, N., Millanes-Romero, A., Pascual-Reguant, L., Morey, L., Diaz, V.M., Lorenz-Fonfria, V., Gutierrez-Gallego, R., Jeronimo, C., Iturbide, A. et al. (2016) Lysyl oxidase-like 2 (LOXL2) oxidizes trimethylated lysine 4 in histone H3. The FEBS journal, 283, 4263–4273.

4. Iturbide, A., Garcia de Herreros, A. and Peiro, S. (2015) A new role for LOX and LOXL2 proteins in transcription regulation. The FEBS journal, 282, 1768–1773.

5. Iturbide, A., Pascual-Reguant, L., Fargas, L., Cebria, J.P., Alsina, B., Garcia de Herreros, A. and Peiro, S. (2015) LOXL2 Oxidizes Methylated TAF10 and Controls TFIID-Dependent Genes during Neural Progenitor Differentiation. Molecular cell, 58, 755–766.

6. Millanes-Romero, A., Herranz, N., Perrera, V., Iturbide, A., Loubat-Casanovas, J., Gil, J., Jenuwein, T., Garcia de Herreros, A. and Peiro, S. (2013) Regulation of heterochromatin transcription by Snail1/LOXL2 during epithelial-to-mesenchymal transition. Molecular cell, 52, 746–757.

7. Barker, H.E., Cox, T.R. and Erler, J.T. (2012) The rationale for targeting the LOX family in cancer. Nature reviews. Cancer, 12, 540–552.

8. Cano, A., Santamaria, P.G. and Moreno-Bueno, G. (2012) LOXL2 in epithelial cell plasticity and tumor progression. Future Oncol, 8, 1095–1108.

9. Ahn, S.G., Dong, S.M., Oshima, A., Kim, W.H., Lee, H.M., Lee, S.A., Kwon, S.H., Lee, J.H., Lee, J.M., Jeong, J. et al. (2013) LOXL2 expression is associated with invasiveness and negatively influences survival in breast cancer patients. Breast Cancer Res Treat, 141, 89–99.

10. Almendro, V., Cheng, Y.K., Randles, A., Itzkovitz, S., Marusyk, A., Ametller, E., Gonzalez-Farre, X., Munoz, M., Russnes, H.G., Helland, A. et al. (2014) Inference of tumor evolution during chemotherapy by computational modeling and in situ analysis of genetic and phenotypic cellular diversity. Cell Rep, 6, 514–527.

11. De Craene, B. and Berx, G. (2013) Regulatory networks defining EMT during cancer initiation and progression. Nat Rev Cancer, 13, 97–110.

12. Escriva, M., Peiro, S., Herranz, N., Villagrasa, P., Dave, N., Montserrat-Sentis, B., Murray, S.A., Franci, C., Gridley, T., Virtanen, I. et al. (2008) Repression of PTEN phosphatase by Snail1 transcriptional factor during gamma radiation-induced apoptosis. Mol Cell Biol, 28, 1528–1540.

13. Vega, S., Morales, A.V., Ocana, O.H., Valdes, F., Fabregat, I. and Nieto, M.A. (2004) Snail blocks the cell cycle and confers resistance to cell death. Genes Dev, 18, 1131–1143.

14. McDonald, O.G., Wu, H., Timp, W., Doi, A. and Feinberg, A.P. (2011) Genome-scale epigenetic reprogramming during epithelial-to-mesenchymal transition. Nat Struct Mol Biol, 18, 867–874.

15. Peinado, H., Moreno-Bueno, G., Hardisson, D., Perez-Gomez, E., Santos, V., Mendiola, M., de Diego, J.I., Nistal, M., Quintanilla, M., Portillo, F. et al. (2008) Lysyl oxidase-like 2 as a new poor prognosis marker of squamous cell carcinomas. Cancer Res, 68, 4541–4550.

16. Herranz, N., Pasini, D., Diaz, V.M., Franci, C., Gutierrez, A., Dave, N., Escriva, M., Hernandez-Munoz, I., Di Croce, L., Helin, K. et al. (2008) Polycomb complex 2 is required for E-cadherin repression by the Snail1 transcription factor. Mol Cell Biol, 28, 4772–4781.

17. Gursoy-Yuzugullu, O., House, N. and Price, B.D. (2016) Patching Broken DNA: Nucleosome Dynamics and the Repair of DNA Breaks. J Mol Biol, 428, 1846–1860.

18. Ziv, Y., Bielopolski, D., Galanty, Y., Lukas, C., Taya, Y., Schultz, D.C., Lukas, J., Bekker-Jensen, S., Bartek, J. and Shiloh, Y. (2006) Chromatin relaxation in response to DNA double-strand breaks is modulated by a novel ATM- and KAP-1 dependent pathway. Nature cell biology, 8, 870– 876.

19. Burgess, R.C., Burman, B., Kruhlak, M.J. and Misteli, T. (2014) Activation of DNA damage response signaling by condensed chromatin. Cell Rep, 9, 1703–1717.

20. Ayrapetov, M.K., Gursoy-Yuzugullu, O., Xu, C., Xu, Y. and Price, B.D. (2014) DNA double-strand breaks promote methylation of histone H3 on lysine 9 and transient formation of repressive chromatin. Proc Natl Acad Sci U S A, 111, 9169–9174.

21. Bonilla, C.Y., Melo, J.A. and Toczyski, D.P. (2008) Colocalization of sensors is sufficient to activate the DNA damage checkpoint in the absence of damage. Molecular cell, 30, 267–276.

22. Khurana, S., Kruhlak, M.J., Kim, J., Tran, A.D., Liu, J., Nyswaner, K., Shi, L., Jailwala, P., Sung, M.H., Hakim, O. et al. (2014) A macrohistone variant links dynamic chromatin compaction to BRCA1-dependent genome maintenance. Cell Rep, 8, 1049–1062.

23. Hodgkinson, A., Chen, Y. and Eyre-Walker, A. (2012) The large-scale distribution of somatic mutations in cancer genomes. Human mutation, 33, 136–143.

24. Schuster-Bockler, B. and Lehner, B. (2012) Chromatin organization is a major influence on regional mutation rates in human cancer cells. Nature, 488, 504–507.

25. Supek, F. and Lehner, B. (2015) Differential DNA mismatch repair underlies mutation rate variation across the human genome. Nature, 521, 81–84.

26. Holliday, D.L. and Speirs, V. (2011) Choosing the right cell line for breast cancer research. Breast cancer research: BCR, 13, 215.

27. Zhang, Y., Liu, T., Meyer, C.A., Eeckhoute, J., Johnson, D.S., Bernstein, B.E., Nusbaum, C., Myers, R.M., Brown, M., Li, W. et al. (2008) Model-based analysis of ChIP-Seq (MACS). Genome Biol, 9, R137.

28. Ernst, J. and Kellis, M. (2012) ChromHMM: automating chromatin-state discovery and characterization. Nat Methods, 9, 215–216.

29. Buenrostro, J.D., Giresi, P.G., Zaba, L.C., Chang, H.Y. and Greenleaf, W.J. (2013) Transposition of native chromatin for fast and sensitive epigenomic profiling of open chromatin, DNA-binding proteins and nucleosome position. Nat Methods, 10, 1213–1218.

30. Tsompana, M. and Buck, M.J. (2014) Chromatin accessibility: a window into the genome. Epigenetics Chromatin, 7, 33.

31. Burgess, R.C. and Misteli, T. (2015) Not All DDRs Are Created Equal: Non-Canonical DNA Damage Responses. Cell, 162, 944–947.

32. Bayona-Feliu, A., Casas-Lamesa, A., Reina, O., Bernues, J. and Azorin, F. (2017) Linker histone H1 prevents R-loop accumulation and genome instability in heterochromatin. Nature communications, 8, 283.

33. Tuduri, S., Crabbe, L., Conti, C., Tourriere, H., Holtgreve-Grez, H., Jauch, A., Pantesco, V., De Vos, J., Thomas, A., Theillet, C. et al. (2009) Topoisomerase I suppresses genomic instability by preventing interference between replication and transcription. Nat Cell Biol, 11, 1315– 1324.

34. Garcia-Rubio, M.L., Perez-Calero, C., Barroso, S.I., Tumini, E., Herrera-Moyano, E., Rosado, I.V. and Aguilera, A. (2015) The Fanconi Anemia Pathway Protects Genome Integrity from R-loops. PLoS Genet, 11, e1005674.

35. Gan, W., Guan, Z., Liu, J., Gui, T., Shen, K., Manley, J.L. and Li, X. (2011) R-loop-mediated genomic instability is caused by impairment of replication fork progression. Genes Dev, 25, 2041–2056.

36. Bakkenist, C.J. and Kastan, M.B. (2003) DNA damage activates ATM through intermolecular autophosphorylation and dimer dissociation. Nature, 421, 499–506.

37. Kaidi, A. and Jackson, S.P. (2013) KAT5 tyrosine phosphorylation couples chromatin sensing to ATM signalling. Nature, 498, 70–74.

38. Yoshida, M., Kijima, M., Akita, M. and Beppu, T. (1990) Potent and specific inhibition of mammalian histone deacetylase both in vivo and in vitro by trichostatin A. J Biol Chem, 265, 17174–17179.

39. Toth, K.F., Knoch, T.A., Wachsmuth, M., Frank-Stohr, M., Stohr, M., Bacher, C.P., Muller, G. and Rippe, K. (2004) Trichostatin A-induced histone acetylation causes decondensation of interphase chromatin. J Cell Sci, 117, 4277–4287.

40. Peinado, H., Del Carmen Iglesias-de la Cruz, M., Olmeda, D., Csiszar, K., Fong, K.S., Vega, S., Nieto, M.A., Cano, A. and Portillo, F. (2005) A molecular role for lysyl oxidase-like 2 enzyme in snail regulation and tumor progression. EMBO J, 24, 3446–3458.

41. Schietke, R., Warnecke, C., Wacker, I., Schodel, J., Mole, D.R., Campean, V., Amann, K., Goppelt-Struebe, M., Behrens, J., Eckardt, K.U. et al. The Lysyl Oxidases LOX and LOXL2 Are Necessary and Sufficient to Repress E-cadherin in Hypoxia: INSIGHTS INTO CELLULAR TRANSFORMATION PROCESSES MEDIATED BY HIF-1. J Biol Chem, 285, 6658–6669.

42. Voloshenyuk, T.G., Landesman, E.S., Khoutorova, E., Hart, A.D. and Gardner, J.D. (2011) Induction of cardiac fibroblast lysyl oxidase by TGF-beta1 requires PI3K/Akt, Smad3, and MAPK signaling. Cytokine, 55, 90–97.

43. Fischer, K.R., Durrans, A., Lee, S., Sheng, J., Li, F., Wong, S.T., Choi, H., El Rayes, T., Ryu, S., Troeger, J. et al. (2015) Epithelial-to-mesenchymal transition is not required for lung metastasis but contributes to chemoresistance. Nature.

44. Zheng, X., Carstens, J.L., Kim, J., Scheible, M., Kaye, J., Sugimoto, H., Wu, C.C., LeBleu, V.S. and Kalluri, R. (2015) Epithelial-to-mesenchymal transition is dispensable for metastasis but induces chemoresistance in pancreatic cancer. Nature.

45. Baldeyron, C., Soria, G., Roche, D., Cook, A.J. and Almouzni, G. (2011) HP1alpha recruitment to DNA damage by p150CAF-1 promotes homologous recombination repair. J Cell Biol, 193, 81–95.

46. Soria, G., Polo, S.E. and Almouzni, G. (2012) Prime, repair, restore: the active role of chromatin in the DNA damage response. Molecular cell, 46, 722–734.

47. Murga, M., Jaco, I., Fan, Y., Soria, R., Martinez-Pastor, B., Cuadrado, M., Yang, S.M., Blasco, M.A., Skoultchi, A.I. and Fernandez-Capetillo, O. (2007) Global chromatin compaction limits the strength of the DNA damage response. The Journal of cell biology, 178, 1101–1108.

48. Goodarzi, A.A., Noon, A.T., Deckbar, D., Ziv, Y., Shiloh, Y., Lobrich, M. and Jeggo, P.A. (2008) ATM signaling facilitates repair of DNA double-strand breaks associated with heterochromatin. Molecular cell, 31, 167–177.

49. Fong, S.F., Dietzsch, E., Fong, K.S., Hollosi, P., Asuncion, L., He, Q., Parker, M.I. and Csiszar, K. (2007) Lysyl oxidase-like 2 expression is increased in colon and esophageal tumors and associated with less differentiated colon tumors. Genes Chromosomes Cancer, 46, 644–655.

50. Moreno-Bueno, G., Salvador, F., Martin, A., Floristan, A., Cuevas, E.P., Santos, V., Montes, A., Morales, S., Castilla, M.A., Rojo-Sebastian, A. et al. (2011) Lysyl oxidase-like 2 (LOXL2), a new regulator of cell polarity required for metastatic dissemination of basal-like breast carcinomas. EMBO molecular medicine, 3, 528–544.

51. Torres, S., Garcia-Palmero, I., Herrera, M., Bartolome, R.A., Pena, C., Fernandez-Acenero, M.J., Padilla, G., Pelaez-Garcia, A., Lopez-Lucendo, M., Rodriguez-Merlo, R. et al. (2015) LOXL2 Is Highly Expressed in Cancer-Associated Fibroblasts and Associates to Poor Colon Cancer Survival. Clinical cancer research: an official journal of the American Association for Cancer Research, 21, 4892–4902.

52. Wong, C.C., Tse, A.P., Huang, Y.P., Zhu, Y.T., Chiu, D.K., Lai, R.K., Au, S.L., Kai, A.K., Lee, J.M., Wei, L.L. et al. (2014) Lysyl oxidase-like 2 is critical to tumor microenvironment and metastatic niche formation in hepatocellular carcinoma. Hepatology, 60, 1645–1658.

53. Martin, A., Salvador, F., Moreno-Bueno, G., Floristan, A., Ruiz-Herguido, C., Cuevas, E.P., Morales, S., Santos, V., Csiszar, K., Dubus, P. et al. (2015) Lysyl oxidase-like 2 represses Notch1 expression in the skin to promote squamous cell carcinoma progression. The EMBO journal, 34, 1090–1109.

54. Bao, S., Wu, Q., McLendon, R.E., Hao, Y., Shi, Q., Hjelmeland, A.B., Dewhirst, M.W., Bigner, D.D. and Rich, J.N. (2006) Glioma stem cells promote radioresistance by preferential activation of the DNA damage response. Nature, 444, 756–760.

55. Wu, M., Wang, P.F., Lee, J.S., Martin-Brown, S., Florens, L., Washburn, M. and Shilatifard, A. (2008) Molecular regulation of H3K4 trimethylation by Wdr82, a component of human Set1/COMPASS. Mol Cell Biol, 28, 7337–7344.

56. Guelen, L., Pagie, L., Brasset, E., Meuleman, W., Faza, M.B., Talhout, W., Eussen, B.H., de Klein, A., Wessels, L., de Laat, W. et al. (2008) Domain organization of human chromosomes revealed by mapping of nuclear lamina interactions. Nature, 453, 948–951.

57. Langmead, B., Trapnell, C., Pop, M. and Salzberg, S.L. (2009) Ultrafast and memory-efficient alignment of short DNA sequences to the human genome. Genome Biol, 10, R25.

58. Kim, D., Pertea, G., Trapnell, C., Pimentel, H., Kelley, R. and Salzberg, S.L. (2013) TopHat2: accurate alignment of transcriptomes in the presence of insertions, deletions and gene fusions. Genome Biol, 14, R36.

59. Trapnell, C., Roberts, A., Goff, L., Pertea, G., Kim, D., Kelley, D.R., Pimentel, H., Salzberg, S.L., Rinn, J.L. and Pachter, L. (2012) Differential gene and transcript expression analysis of RNA-seq experiments with TopHat and Cufflinks. Nat Protoc, 7, 562–578.

60. Lerat, E., Fablet, M., Modolo, L., Lopez-Maestre, H. and Vieira, C. (2017) TEtools facilitates big data expression analysis of transposable elements and reveals an antagonism between their activity and that of piRNA genes. Nucleic Acids Res, 45, e17.

61. Tarazona, S., Furio-Tari, P., Turra, D., Pietro, A.D., Nueda, M.J., Ferrer, A. and Conesa, A. (2015) Data quality aware analysis of differential expression in RNA-seq with NOISeq R/Bioc package. Nucleic Acids Res, 43, e140.

62. Quinlan, A.R. and Hall, I.M. (2010) BEDTools: a flexible suite of utilities for comparing genomic features. Bioinformatics, 26, 841–842.

